# DirectMS1: MS/MS-free identification of 1000 proteins of cellular proteomes in 5 minutes

**DOI:** 10.1101/756213

**Authors:** Mark V. Ivanov, Julia A. Bubis, Vladimir Gorshkov, Irina A. Tarasova, Lev I. Levitsky, Anna A. Lobas, Elizaveta M. Solovyeva, Marina L. Pridatchenko, Frank Kjeldsen, Mikhail V. Gorshkov

## Abstract

Proteome characterization relies heavily on tandem mass spectrometry (MS/MS) and is thus associated with instrumentation complexity, lengthy analysis time, and limited duty-cycle. It was always tempting to implement approaches which do not require MS/MS, yet, they were constantly failing in achieving meaningful depth of quantitative proteome coverage within short experimental times, which is particular important for clinical or biomarker discovery applications. Here, we report on the first successful attempt to develop a truly MS/MS-free and label-free method for bottom-up proteomics. We demonstrate identification of 1000 protein groups for a standard HeLa cell line digest using 5-minute LC gradients. The amount of loaded sample was varied in a range from 1 ng to 500 ng, and the method demonstrated 10-fold higher sensitivity compared with the standard MS/MS-based approach. Due to significantly higher sequence coverage obtained by the developed method, it outperforms all popular MS/MS-based label-free quantitation approaches.

---

Advances in mass-spectrometry-based proteomic technologies resulted in dramatically increased depth, throughput, and sensitivity of proteome coverage. Up to 10,000 proteins can be identified in an 100 minute analysis of human cell proteomes using state-of-the-art high-resolution Orbitrap mass spectrometry^1^. Recently, the notable trend in LC-MS technology developments has been toward increasing the throughput of the proteome-wide analysis, while preserving the quantitation accuracy^2,3^. However, these achievements rely heavily on the use of tandem mass spectrometry (MS/MS), which includes sequential isolation of eluting peptides followed by their fragmentation. While being a crucial and seemingly the only source of sequence-specific information about the peptides, MS/MS brings a number of well-known challenges. Due to the limited both the speed of the mass analyzer (which is almost exclusively Orbitrap FTMS^4^) and the peak capacity of the separation system multiplied by the proteome digest complexity, the fragmentation spectra are only produced for a fraction of all ionized peptides from the sample. Any decrease in the analysis time, e.g. by using shorter LC gradients, reduces this fraction even more^5^. As a result, only a few peptides are identified for each protein. This undermines quantitation accuracy and leads to a bias toward identification of larger and/or more abundant proteins. Indeed, protein quantitation is a crucial value of proteomics in clinical and/or biomarker discovery applications. Among the available approaches, the most popular and more suitable for high-throughput proteome-wide analyses are label-free quantitation methods (LFQ)^6–9^. However, these methods strongly depend on the protein sequence coverage, which typically reduces to one or two identified peptides for short LC gradients.

The above-mentioned problems are further aggravated by the complexity of the whole proteome digests and limited peak capacity of existing LC systems. The latter results in appearance of co-eluted peptides originating from different proteins observed in the same *m/z* isolation window. Being fragmented together, these peptides produce the so-called chimeric fragmentation spectra, which can be wrongly attributed to a peptide not present in the sample^10^ and further undermine the performance of database search engines^11^. Again, the obvious solution to the latter problem is using longer gradients for separation combined with LC columns of significantly extended lengths and extensive sample pre-fractionation, thus, further extending the analysis time due to prolonged column equilibration, etc^12^.

In general, the above factors, associated mostly with the use of MS/MS, undermine the utility of proteome analysis in biomarker discovery and in the clinical environment, in which hundreds of samples have to be quantified proteome-wide within days, if not hours^13^. These arguments advocate for the need for even a faster proteome analysis time down, preferably, to a minute time scale. One of the obvious routes to this goal is reducing the time spent acquiring MS/MS spectra, or removing it completely from the experimental pipeline. For years, it was tempting to implement approaches which do not require MS/MS for protein identification and quantitation, starting from peptide mass fingerprinting in the earlier days of bottom-up proteomics, and to the numerous recent efforts based on utilization of complementary sequence-specific peptide properties or labeling techniques^14,15^. However, most of these approaches fail to achieve the meaningful depth of quantitative proteome coverage within short experimental times. Many of the recent efforts were focused on hybrid experimental pipelines, in which MS/MS-based identification is combined with MS1-based quantitation^3,16^.

Progress in mass spectrometry allows the masses of biomolecular ions to be measured with sub-ppm precision. However, this accuracy is still far from being sequence-specific for peptides^17^, the deficiency countered by MS/MS^18^. Recent studies have shown that retention times are sequence specific in case of peptides and proteins and may potentially replace MS/MS^19^. This intrinsic value of the chromatography data was also extensively explored, mainly, to support peptide identification and validation by reducing the search space or helping identification of the features in MS1 spectra^14,20^. One of the main obstacles associated with the use of chromatography for implementing true MS1-based proteomics workflows was its relatively low accuracy in prediction of retention times for peptides, which is still much lower compared with the LC’s experimental precision and reproducibility^19^. In other words, exact masses measured with sub-ppm accuracy and retention times alone were not enough to identify a tryptic peptide from a database unambiguously. More complementary data seems to be needed to remove this ambiguity by adding more dimensions to the MS1-based search space. A number of efforts in this direction have been reported recently, including the use of partial peptide sequence degradation based on Edman’s reaction^21^ and isotopic labeling^15^. A truly MS1-based workflow called *ms1 searchpy* was introduced recently based on 2D (retention time, mass) search space for peptide identification and the sequence-specific orthogonality of parallel digestion using different enzymes^22^. The method demonstrated that MS1-only proteome-wide analysis is possible; yet, the use of different enzymes requires further efforts in finding the enzymes of digestion specificity and reproducibility comparable with trypsin.

In this work we have integrated *ms 1 searchpy* algorithm with a number of utilities for LC-based peptide feature detection, retention time prediction, and protein quantitation. This novel integrated software allows for the first time to break the 1000 protein identification barrier for MS1-only quantitative proteome analysis based on 2D (retention time, mass) search space. Moreover, this level of identification efficiency was achieved in 5-minute LC-MS analysis.

## RESULTS

### DirectMS1 method: the workflow

The MS1-only workflow, which we call DirectMS1, is shown in **Figure 1**. In this method, a high resolution mass analyzer is used to acquire MS1 spectra at the highest possible speed and mass measurement accuracy. For example, currently available Orbitrap FTMS mass analyzers are capable of acquiring up to 4 MS spectra per second at the resolution of 120,000 at *m/z* 200. This acquisition rate allows using ultra-short HPLC gradients for separating the whole proteome digests. Indeed, the chromatographic peaks for peptides are typically 3 to 5 seconds wide for gradients of a few minutes in duration, thus allowing acquisition of up to 20 mass spectra per each eluting peptide. This quantity is important for subsequent peptide feature detection.

**Figure 1.**
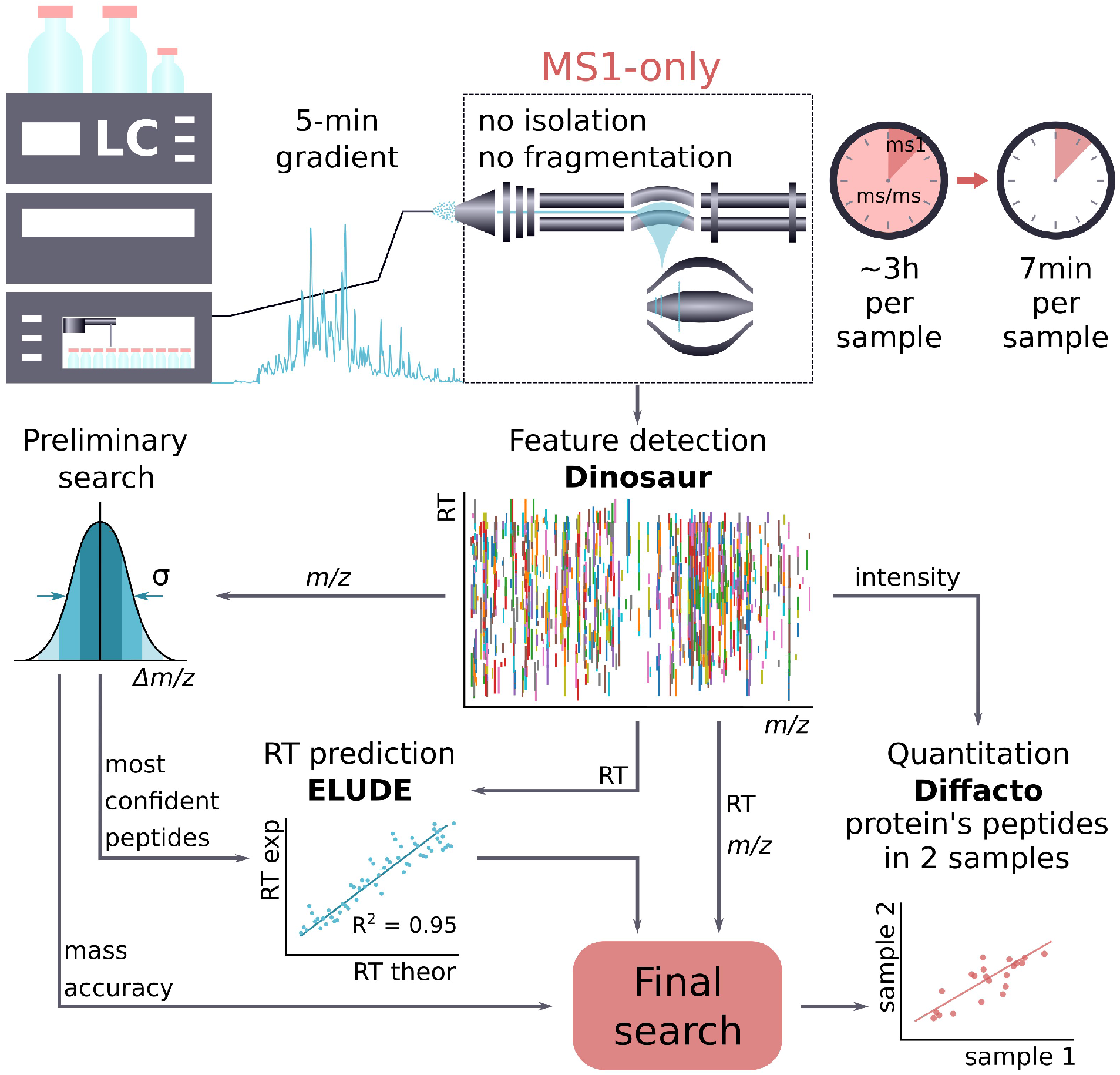
DirectMS1 workflow. Protein identification and quantitation are done using LC-MS1 data.

Protein identification is implemented in *ms 1 searchpy* software described earlier^22^. For the latest implementation, the algorithm was significantly modified to improve its efficiency (by both the calculation speed and the number of identified proteins), *ms1searchpy* in turn integrates three software utilities: *Dinosaur*^24^ for deisotoping of mass spectra and peptide feature detection, *ELUDE*^25^ for machine-learning-based peptide retention time prediction, and *Diffacto*^26^ for label-free MS1-based protein quantitation, *ms1 searchpy* itself matches peptide features found in the MS1 spectra to theoretical peptides using measured *m/z* and retention time (RT) values, and uses these matches to calculate the protein scores based on the binomial distribution model, *ms1searchpy* is open-source and freely available at https://bitbucket.org/markmipt/ms1searchpy under Apache 2.0 license. More details on the *ms1searchpy* software can be found in the Supporting materials.

### Proteome coverage using DirectMS1

The efficiency of proteome analysis is characterized by the number of identified peptide spectrum matches and/or proteins at the specified level of false discovery rate (FDR). In this study, we focused on the number of protein groups identified at 1% FDR and compared the results with the MS/MS-based approach using the same chromatographic separation time. To maximize the efficiency of MS/MS analysis, we used *IdentiPy* search engine^27^, which features built-in resolution of potentially chimeric spectra, and the recently introduced machine learning algorithm *Scavager* for postsearch validation of peptide and protein identifications^28^. The results of this “*IdentiPy* + *Scavager*” combination are presented in the manuscript as “MS2 data”. **Figure 2a** shows the dependence of DirectMS1 efficiency on the MS1 mass resolving power. In these experiments we loaded 200 ug of HeLa digest and ran 4.8-minute HPLC gradients with the total HPLC method time of 7.3 minutes. Mass resolving power is one of the key factors affecting the efficiency of the method, as it relies on detecting peptide features in the spectra which have to be resolved under the ultra-short separation conditions in the first place. It also affects the mass measurement accuracy. On the other hand, working with the highest possible resolution settings can be detrimental because of decreasing number of acquired spectra within the peptide elution time. For the three technical replicates we were able to identify 923 protein groups at 1% FDR on average. These results were achieved at 120k mass resolution. More details on the mass measurement accuracy, number of detectable features, and the number of MS1 scans for different mass resolution settings are shown in **Fig. 2b-d**. Note that the number of detectable features is almost the same for 120k and 240k mass resolution settings, but longer acquisition time per spectrum for the latter results in lower number of MS1 scans and identified protein groups. Another factor contributing negatively to the method’s efficiency at high resolving powers is the higher level of noise due to longer acquisition time, which is translated into false positive peptide features. The standard MS/MS-based method yields less than 500 protein groups for the same chromatographic separation time. Expectedly, its efficiency decreases with increasing mass resolution as it depends on the time available for acquiring MS/MS spectra for as much precursors as possible. Also, the problem of co-eluting peptides of close *m/z* and massive appearance of chimeric spectra becomes acute for this method for short separation times.

**Figure 2.**
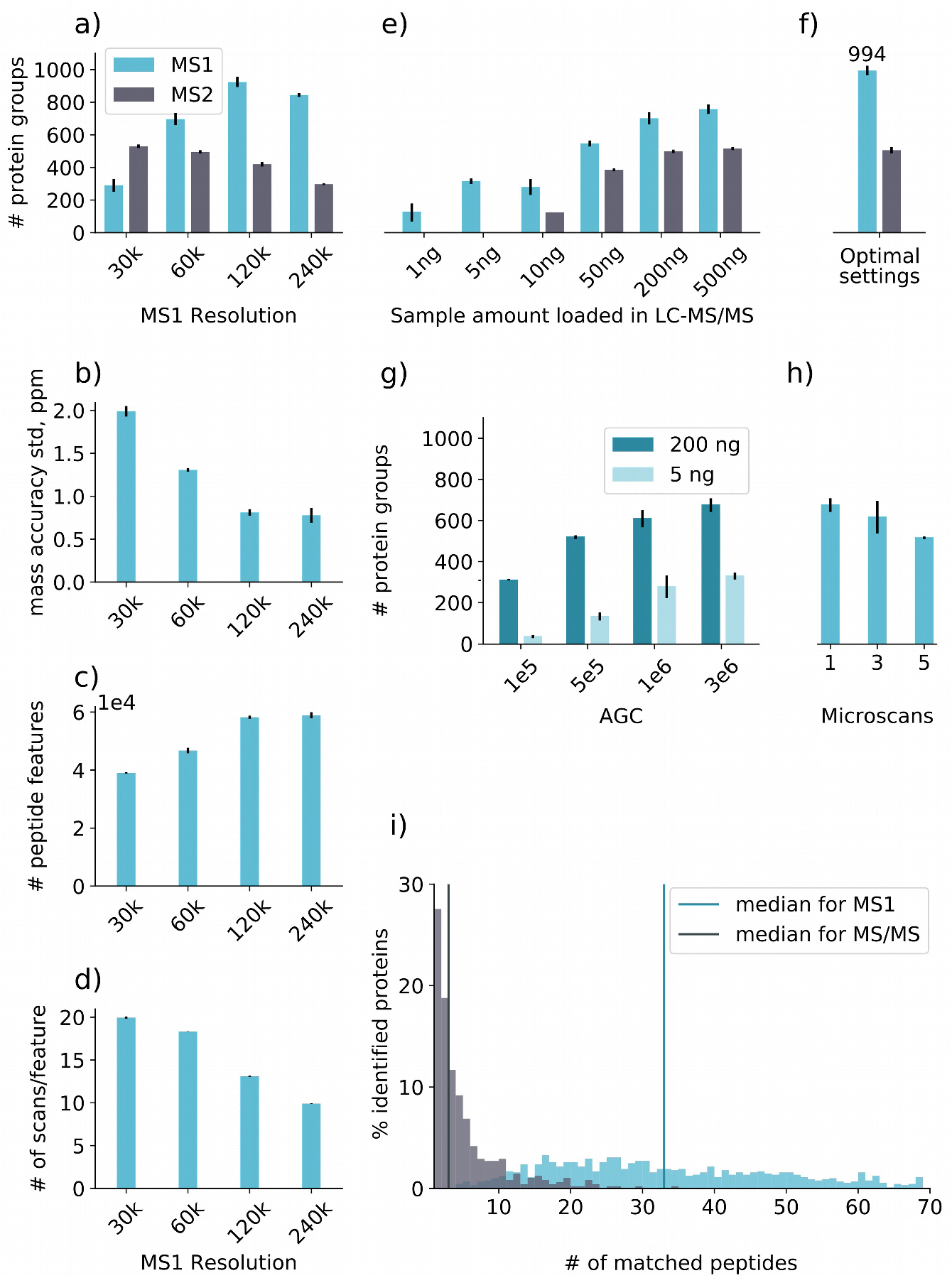
Proteome analysis of HeLa cell line using 5-minute HPLC separation gradient, different sample loads and mass resolution settings, as well as the comparison of MS/MS-based and DirectMS1 methods: **a** – number of identified protein groups for different MS1 resolution settings; **b** – mass measurement accuracy for peptides in MS1 spectra; **c** – number of detected peptide features; **d** – average number of scans per peptide feature detected; **e** – number of identified protein groups for different sample amount loaded; **f** – results for 500 ng loaded HeLa amount and 120k MS1 resolution; **g** – results for the analysis of different amounts of loaded samples using different AGC settings; **h** – results for acquisition of MS1 spectra at varying number of summed microscans; **i** – protein sequence coverage (results are shown for 120k mass resolution). The identification results are shown for the average values obtained for 3 technical replicates (except panel *i*). Results are shown for 1% protein group FDR.

Next, the amount of HeLa sample at the protein level loaded for the analysis was varied from 1 to 500 ng (**Fig. 2e**). Mass resolution settings were 60k in these experiments, as an efficiency trade-off between the two methods according to the results shown in **Fig. 2a**. We could not identify a single protein group at 1% FDR using the MS/MS-based method when the amount of loaded sample was 10 ng and less, while DirectMS1 gave more than 100 protein groups even for 1 ng of loaded sample. Indeed, the sensitivity of MS1 acquisition is higher compared with tandem mass spectrometry *a priori* (for the price of specificity, of course). Thus, in both methods the peptide features are detected in MS1 spectra by the analyzer. However, the amount of precursor ions that can be accumulated within a reasonable time to perform fragmentation becomes too small for obtaining MS/MS spectra of sufficient quality. To further explore the issue with the analysis sensitivity the target amount of ions accumulated in the external radio-frequency (rf) ion trap prior to injection into the high-resolution mass analyzer (the so-called Automatic Gain Control value – AGC) was varied from 10^5^ to 3*10^6^ charges for two different sample loads, 5 ng and 200 ng. The results are shown in **Fig. 2g**. For both loads the largest numbers of identified protein groups were obtained for the highest AGC value, contrary to some reasonable expectation. Indeed, decreasing the AGC leads to improvement in mass measurement accuracy (**Supplementary Figure S3**) because of reduced space charge effect^29^. Moreover, this reduction helps avoiding the ion coalescence problem in MS1 spectra,^30,31^ which may be especially pronounced under the conditions of ultra-short separations. Thus, the efficiency of DirectMS1 method should potentially increase with lower AGC. However, the number of detectable peptide features has dramatically dropped, resulting in a lower number of identified protein groups. We attribute this observation to the decreasing signal-to-noise ratio for the observed peaks, which hinders feature detection. Summing several scans for each MS1 acquisition proved to be ineffective for the current version of data processing software, as shown in **Fig. 2h**. Single scans provide a higher number of identifications in spite of the seemingly obvious decrease in signal-to-noise ratios for the peaks in the spectra compared with 3 and 5 scan summations. We attribute this effect to the *Dinosaur* software used for peak picking and deisotoping, which has built-in scan averaging, and the summation of several scans simply leads to a lower MS1 acquisition rate.

Upon optimization of all experimental parameters affecting the proteome analysis efficiency in ultra-short separations, a direct comparison of both DirectMS1 and standard MS/MS-based methods was performed. The complete set of optimized parameters is provided in the Method section below. The mass resolution was set to 120k at *m/z* 200, the amount of HeLa digest loaded was 500 ng, 1 microscan with AGC target of 3*10^6^ was used. The methods delivered identification of 994 (up to 1024 in the single run) and 506 protein groups on average for 3 replicate runs, for DirectMS1 and data-dependent MS/MS method respectively as shown in **Fig. 2f**. Note that the numbers for protein groups identified using DirectMS1 are more conservative compared with the MS/MS-based analysis. The latter reports a protein group even if protein identification is based on a single unique peptide based on high sequence specificity of tandem mass spectrometry. Yet, such a “one-hit-wonder” protein would not pass the identification threshold in DirectMS1 approach. All shared peptides will be scored only once for the most confident protein, and the rest of proteins will be scored based on unique peptides only. The minimal number of peptides typically required for successful protein identification in DirectMS1 is three, as shown in **Fig. 2i**, and the median number of peptides identified per protein is 3 and 32 for MS/MS and DirectMS1, respectively. This is an important difference between the methods: DirectMS1 provides significantly higher sequence coverage for identified proteins (even compared with the hour-long MS/MS-based proteome analyses), which can be beneficial for quantitation.

### Quantitation

Protein quantitation provides an additional important dimension of proteome analysis. Label-free quantitation (LFQ) approaches remain among the most popular methods. They are inexpensive, easy to implement, and allow rapid proteome-wide estimation of relative protein concentrations across multiple samples. DirectMS1 was compared with three different LFQ approaches applicable to MS/MS-based analysis, including: *MaxQuant* LFQ, which is probably one of the most widely used workflows in shotgun proteomics; *IdentiPy* + NSAF^9^; and *IdentiPy* + *Diffacto*. NSAF was selected as one of the best LFQ algorithms as shown previously^8^. Diffacto was utilized for DirectMS1 method. This algorithm uses the identified peptide ion peak intensities in MS1 spectra and applies factor analysis to extract covariation between peptide abundances, which in turn provides estimates of protein abundances. Here, for the analysis of MS/MS-based data, we extended the *IdentiPy* search engine to allow running the *Dinosaur* software to find peptide features in MS1 spectra followed by the de-multiplexing of chimeric MS/MS spectra in a way similar to the previously described DeMix algorithm^32^. Along the way, the MS1 peaks intensities of peptide ions were also extracted for identified MS/MS spectra and used by *Diffacto*.

Comparison of the methods was performed using six proteins of varying concentrations spiked into the yeast proteome. Details of the analyzed mixtures are shown in **Supplementary Table S1**. The concentration of one of the proteins was always significantly higher than the others to better reflect the dynamic range of the biological samples, in which a few proteins may be present at concentrations exceeding the rest by several orders of magnitude (e.g., human plasma samples). As a result, a mass analyzer spends most of the time acquiring MS/MS spectra of a few highly abundant peptides from a few major proteins, which creates a bias in quantitation of the other proteins. The problem becomes especially important for rapid proteome analyses employing ultra-short separation gradients.

The results obtained here for different LFQ methods were compared using two metrics: number of true/false positives after applying statistical tests and the accuracy of protein concentration measurement, p-values were calculated using either *Diffacto* algorithm, or the one-way ANOVA test, and the p-value threshold of 0.05 with Bonferroni correction was chosen as significant.

The accuracy of concentration measurement was compared by plotting ratios of protein concentrations between samples for experimental and calculated values. The metric used for the accuracy was standard quantification error (*SQE*). *SQE* is defined as the root-mean-square error of (logarithmized) calculated concentration ratios of target proteins:

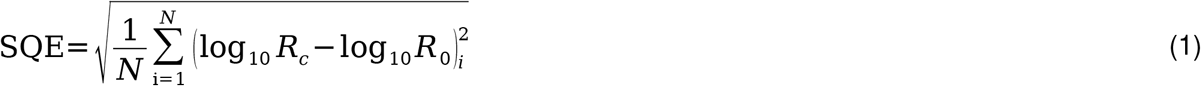

in which *R_c_* is the ratio of calculated abundances for the same protein in two different samples (calculated ratio), *R_0_* is the ratio of actual concentrations of the proteins (“actual” ratio), and *N* is the number of ratios for all proteins. Lower *SQE* values means that a particular method estimates the relative protein concentrations more accurately.

**Fig. 3a-b** shows the results of LFQ accuracy tests. DirectMS1 analysis, which has *Diffacto* algorithm built in, has the average *SQE* of 0.301, and is the most accurate among the methods evaluated. Importantly, this method demonstrates the highest quantitation accuracy for all 5 proteins with significantly altered concentrations. For the other methods, we were able to calculate *SQE* for 3 reported proteins only. MS/MS-based analyses with NSAF, *Diffacto*, and MaxLFQ algorithms have the average *SQEs* of 0.43, 0.747, and 0.742, respectively.

**Figure 3.**
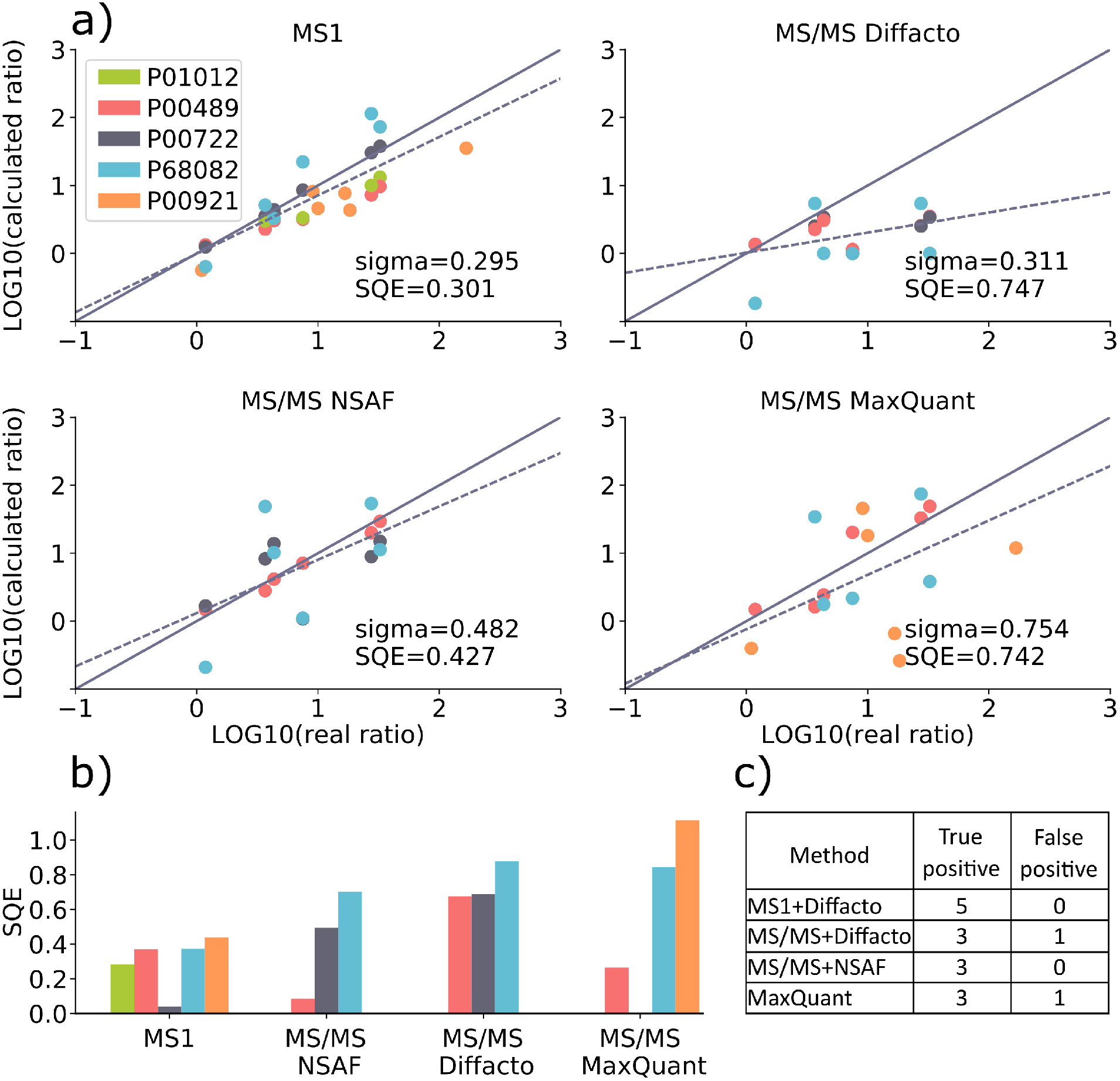
Results of the quantitation analysis using spiked-in mixtures of 6 proteins in yeast proteome: (**a**) protein abundance ratios for experimental and calculated concentrations. Abundances of 5 proteins were compared between 4 samples. For MS/MS based methods only 3 proteins which were reported as significantly changed are shown; (**b**) standard quantification errors (*SQE*) estimated for the evaluated LFQ methods. The lower is the *SQE*, the better is the quantitation accuracy. The missing bars correspond to proteins which have not passed the significance threshold, (**c**) number of proteins which passed 0.05 p-value threshold with Bonferroni correction for DirectMS1 method based on *Diffacto* quantitation algorithm, *IdentiPy* search engine with added *Diffacto* or NSAF quantitation algorithms, and *MaxQuant* with MaxLFQ. Here, the spiked-in proteins are considered true positives and yeast proteins from the background are considered false positives.

For DirectMS1 method, all five significantly altered proteins and none of the yeast proteins have passed the threshold, which was the best results among the methods evaluated, as shown in **Fig. 3c**. For MS/MS+*Diffacto*, MS/MS+NSAF, and *MaxQuant* the results were 3/1, 3/0 and 3/1, respectively.

### Conclusions

We developed a method of whole proteome analysis DirectMS1, which does not employ tandem mass spectrometry. The method allows using ultra-short separation gradients, for which it considerably outperforms traditional MS/MS-based approaches in depth of the proteome coverage, protein quantitation accuracy, and sensitivity. Specifically, we have demonstrated the identification of more than 1000 proteins of the human cell line proteome in 5 minutes, also breaking this pivotal identification number in MS/MS-free proteomics for the first time. The method was also able to identify more than 100 proteins when the amount of loaded HeLa digest sample was 1 ng only. In addition to significantly increased proteome analysis throughput, which is important in clinical proteomics, the research community can benefit from the method by employing a simpler mass spectrometry instrumentation. We expect further improvements in the method’s performance from development of more accurate retention time prediction models and new peak picking algorithms. One limitation is that the method currently does not support PTM studies due to its high sensitivity to the size of the search space.

## ACKNOWLEDGMENTS

This study was supported by the Russian Science Foundation: grant #14-14-00971 for development of *ms1searchpy* and *Identipy* software packages and preliminary evaluation of their efficiencies, #19-74-00123 for development and testing of MS1-based quantitation tools based on Diffacto; the European Research Council (ERC) under the European Union’s Horizon 2020 Research and Innovation Programme (grant agreement No. 646603); and the VILLUM Center for Bioanalytical Sciences at the University of Southern Denmark. Authors also thank Dr. Dmitry S. Karpov for providing yeast cell line samples.

## METHODS

### Samples

Development of the method and its evaluation were performed using Thermo Scientific Pierce™ HeLa Protein Digest Standard (P/N 88328) derived from HeLa S3 cell line. For the quantitation part of the study a wild-type yeast (strain BY4741, Euroscarf, Germany) with 6 spiked-in proteins (manufacturers are listed in **Supplemental Table S1**) of different molecular weights and sequence lengths was used. The concentrations of 5 of these proteins (see Supplemental Table S1) were varied in a range from 1 fM to 100 fM, while the sixth protein, BSA, was spiked into the yeast proteome at the concentration of 500 fM in all mixtures.

### Sample preparation

Yeast cells were handled in the following way: aliquot of approximately 10^7^ cells were resuspended in 100 μL of lysis buffer (0.1 % w/v ProteaseMAX Surfactant (Promega, USA) in 50 mM ammonium bicarbonate and 10 % v/v ACN). The cells were then incubated in a shaker for 1h at 550 rpm at room temperature. Cells were lysed using ultrasonic homogenizer BandelinSonopuls HD2070 (Bandelin Electronic, Berlin, Germany) by sonication for 2 minutes at each 30, 60, 80 % amplitudes on ice. The supernatant was collected after centrifugation at 13 000 rpm for 10 min at room temperature (Centrifuge 5415R; Eppendorf, Hamburg, Germany). Total protein concentration was measured using BCA assay. Protein extracts were reduced in 10 mM DTT at 56 ^<u>o</u>^C for 20 min and alkylated in 10 mM iodoacetamide at room temperature for 30 min in dark. Then, samples were digested overnight at 37 ^<u>o</u>^C using trypsin protease (Sequencing Grade Modified Trypsin, Promega, Madison, WI, USA) added at the ratio of 1:50 w/w. Enzymatic digestion was terminated by the addition of acetic acid (5 % w/v). After the reaction was stopped, the samples were shaken (550 rpm) for 25 min at room temperature followed by centrifugation at 13 000 rpm for 10 min at 20 ^<u>o</u>^C (Centrifuge 5415R; Eppendorf, Germany). Then the supernatant was dried in SpeedVac at 45 ^<u>o</u>^C. Peptides were stored at −80 ^<u>o</u>^C until the LC-MS/MS analysis. Before the LC-MS/MS analysis, the samples were desalted using Oasis cartridges for solid phase extraction (Oasis HLB, 1 cc, 10 mg, 30 μm particle size, Waters). Then, the peptide concentration for each sample was measured using the BCA assay. Six spike proteins were mixed according to ratios in Table S1, then mixes were reduced in 10 mM DTT at 56 ^<u>o</u>^C for 20 min and alkylated in 10 mM iodoacetamide at room temperature for 30 min in the dark. Proteins were digested overnight using trypsin protease (Sequencing Grade Modified Trypsin, Promega, Madison, WI, USA) in 1:50 w/w ratio. Then, protein digests were spiked to 1 ug yeast for one LC-MS/MS injection.

### LC-MS/MS methods

LC-MS/MS analysis was performed using Orbitrap Q Exactive HF mass spectrometer (Thermo Fisher Scientific, San Jose, CA, USA) coupled with UltiMate 3000 LC system (Thermo Fisher Scientific, Germering, Germany). Mass spectrometry measurements were performed either in data-dependent acquisition (DDA) mode with “Top15” setting for MS/MS spectra, or in MS1-only mode of acquisition. By default, the Full MS scans were acquired from *m/z* 375 to 1500 at a resolution of 60k at *m/z* 200 with a target of 3·10^6^ charges for the automated gain control (AGC), 1 microscan and 200 ms maximum injection time. For higher-energy collision-induced dissociation (HCD) MS/MS scans, the normalized collision energy was set to 30, the resolution was 15k at *m/z* 200. Precursor ions were isolated in a 1.4 Th window and accumulated for a maximum of 30 ms or until the AGC target of 2·10^5^ ions was reached. Precursors of charge states from 2+ to 7+ were scheduled for fragmentation. Previously targeted precursors were dynamically excluded from fragmentation for 4 s. 200 ng of HeLa and 1000 ng of yeast digests were loaded on column by default. **Supplementary Table S2** contains list of all raw files with brief description. Short gradient LC method was adopted from the following technical note provided by the vendor (https://assets.thermofisher.com/TFS-Assets/CMD/Technical-Notes/tn-72827-lc-ms-tandem-capillary-flow-tn72827-en.pdf) with minor changes. Trap column μ-Precolumn C18 PepMap100 (5 μm, 300 μm, i.d. 5 mm, 100 Å) (Thermo Fisher Scientific, USA) and self-packed analytical column (Inertsil 3 μm, 75 μm i.d., 15 cm length) were employed for separation. Mobile phases were as follows: (A) 0.1 % FA in water; (B) 80 % ACN, 0.1 % FA in water. Loading solvent was 0.05 % TFA in water. The gradient was from 5 % to 35 % phase B in 4.8 min at 1.5 μL/min. Total method time was 7.3 min.

### Protein identification

Raw files were converted to mzML format using *ProteoWizzard*^33^ (v. 3.0.5533). MS1 spectra were processed by *ms1 searchpy* (v. 1.1.2) algorithm^22^, *ms1searchpy* uses *Dinosaur*^24^ (v. 1.1.3) for feature detection and *ELUDE*^25^ (v. 3.02.1) for retention time prediction. The parameters for MS1 search engine were the following: 5 ppm precursor mass accuracy, 0 missed cleavages, carbamidomethylation of cysteine as fixed modification, minimal peptide length of six amino acids, 1 to 5 charge states, minimal 2 visible 13C isotopic peaks in the isotopic envelope detected in at least 3 scans were allowed for identifying a peptide feature. *IdentiPy*^27^ (v. 0.2) and *MaxQuant*^34^ (v. 1.6.5.0) search engines were used for MS/MS-based searches. The parameters for MS/MS search engines were the following: 10 ppm precursor mass accuracy, 0.05 Da fragment mass accuracy, 2 missed cleavages, carbamidomethylation of cysteine as fixed modification, minimal peptide length of six amino acids. DeMix algorithm^32^ for chimeric spectra processing was integrated into *IdentiPy* search engine for increasing the search efficiency. *IdentiPy* search result files were postprocessed using *Scavaget*^28^ (v. 0.1.9).

### False discovery rate estimation

For HeLa searches human Swiss-Prot database containing 20193 sequences was used. Yeast database (6621 proteins) including 6 spiked proteins was used for quantitation part of the study. Standard target-decoy strategy^35^ (TDS) was used for FDR estimation. Decoys were generated by shuffling sequences of target proteins using Pyteomics^36^. For additional validation of developed workflow, DirectMS1 method was tested using extended protein database. Yeast protein sequences were combined with the human ones followed by generation of shuffled decoy proteins. Assuming that there are no yeast proteins in the sample, the real level of FDR can be estimated for the results of MS1 searches. Apparently, there was only 1 target yeast protein found among 1021 proteins in the results (FDR 0.1 %), which is close to the expected value of 0.25 %.

