## Supporting Materials for "DirectMS1: MS/MS-free identification of 1000 proteins of cellular proteomes in 5 minutes"

**Description of the workflow**. Detailed workflow of *ms1searchpy* software is shown in **Supplementary Figure S1**. Peptide features are extracted from mzML file at the first step using *Dinosaur* software. Next, *ms1searchpy* matches peptide features to theoretical peptides for target and decoy proteins using user defined mass accuracy, typically 10 ppm, which is enough to account for potential systematic mass shifts. The probability of a peptide to be matched randomly can be calculated using the following equation:

$$p=\frac{k_d}{N_d} \quad (1),$$

in which $k_d$ is the number of matched decoy peptides, and $N_d$ is the total number of decoy peptides for the search space. Then, the protein score, *ProteinScore*, is calculated as follows:

$$\text{ProteinScore}=-\log_{10}\left(\text{Sf}\left(\text{k,N,p}\right)\right) \quad (2),$$

in which $k$ is the number of matched peptides, $N$ is the number of theoretical peptides for a protein, $Sf$ - is the survival function for the binomial distribution.

At the initial step, the protein scores are calculated for all proteins in the database, and the list of top-scoring proteins is constructed and filtered to 1% FDR using the target-decoy approach.

Typically, this step produces 10 to 100 proteins. For these proteins, the distribution of mass errors is expected to be a sum of normal distribution (true peptide identifications) and uniform distribution (false peptide identifications). ms1searchpy fits the data obtained and estimates the mean and the standard deviation for the normal distribution. The mean +/- 3 standard deviations is used as mass accuracy range in the next step. Here, the peptide matching and protein scores recalculated again, and up to 1000 most reliably identified peptides from proteins filtered to 1% FDR are selected. Reliability is determined as a minimal Z-score for the peptide mass error. These peptides are used for training the retention time prediction model and the estimation of its accuracy. In the current version of DirectMS1 method, ELUDE is used as a model for RT prediction. Alternatively, we have incorporated the less sophisticated retention time prediction additive model, but the method's efficiency decreased by 10 to 20% for the number of identified protein groups (**Supplementary Table S3**). This observation demonstrates the importance of accuracy in retention time prediction for the method's performance. At the final step, the protein scores are recalculated again using +/- 3 Z-score filter for mass and retention time tolerances. At this stage, two important additions to the workflow are implemented (see the pseudo code below). The first one provides protein grouping. All proteins are sorted according to the calculated score values. Thereafter, the best scoring protein is moved into the search results and all peptide features related to this protein are removed before the next iteration ("one feature - one peptide" rule). At the same time, all peptides related to this protein are kept in the memory to prevent the use of shared peptides by the other proteins at the later steps ("one peptide - one protein" rule) (see **Supplementary Figure S2**). At the next step, the probability of a peptide to be matched randomly and all scores for the rest of the proteins are recalculated and the next best scoring protein is also moved into the search results, etc. The above iteration is repeated until FDR of the proteins in the search results exceeds FDR of 10%. The above steps form the list of protein groups ensure that shared features and peptides are counted only for one protein.

*Pseudo code for the iteration procedure in ms1searchpy workflow*

```
--------------------------------------------------------------------------------------
input all_PFMs, proteins_preliminary_scores
initialize final_proteins

# All shared peptides are kept only for high scoring protein among all #alternative proteins for these peptides ("one peptide - one protein" rule):
all_PFMs.remove_duplicates(proteins_preliminary_scores)

thresholds = [100%, 66%, 33%]
for mass_threshold in thresholds:
    for RT_threshold in thresholds:
        # Keep X% best matches by mass error
        filtered_PFMs = filter(all_PFMs, mass_threshold)
        # Keep Y% best matches by RT error
        filtered_PFMs = filter(filtered_PFMs, RT_threshold)

        initialize filtered_proteins
        while FDR(filtered_proteins) <= 10%:
            all_protein_scores = calculate_scores(filtered_PFMs)
            best_protein, best_protein_score = get_best_scored(all_protein_scores)
            filtered_proteins.add(best_protein, best_protein_score)

            # Remove all features which are matched by peptides from the best protein
            best_protein_features = get_features(filtered_PFMs, best_protein)
            filtered_PFMs.remove(best_protein_features)

        #Add rest of the proteins without score recalculating
        for protein in all_protein_scores:
            if protein not in filtered_proteins:
                filtered_proteins.add(protein)

        # Add filtered proteins from current iteration to the final list
        final_proteins.add(filtered_proteins)

# Get average protein score when all iterations are finished
final_proteins = average_score(final_proteins)
return filter_FDR(final_proteins, FDR=1%)
--------------------------------------------------------------------------------------
```

The important part of the method is mass and retention time iterative filtering. By design of the matching procedure, all true and false identifications lie within the ±3Z range. It is expected that the deviation of the calculated peptide masses and retention times from the measured ones are distributed normally and uniformly for the true and false peptide identifications, respectively. Thus, inside the +/-2 Z-score range, there will be 95% of true and 66% of false identifications. Likewise, for the +/-1 Z-score filter, we expect to have 65% and 33% of true and false identifications, respectively. However, there is a trade-off between keeping the maximal number of true identifications and lowering the number of false ones, and there is no a single unique threshold existing here. The iteration procedure is then aimed at calculation of the optimal protein score by looking at the "best" peptides in terms of mass and retention time errors. At the first step of this procedure, all peptides within +/-3 Z-score mass and RT errors range are used for the scoring, yet, taking into account the "one peptide - one protein" rule from the previous step. This rule is crucial to have confidence in that the protein group leaders will not be changed during the iterations. Thereafter, the protein scores are recalculated for all thresholds in combinations of +/-1, 2 and 3 Z-score for mass and RT errors. For example, first iteration will be done by keeping all peptide identifications within +/-2 and +/-3 Z-scores for mass and RT errors, respectively. The last one - for +/-1 and +/-1. After all iterations, we have 9 different scores for each protein, which are averaged and used as the final protein score.

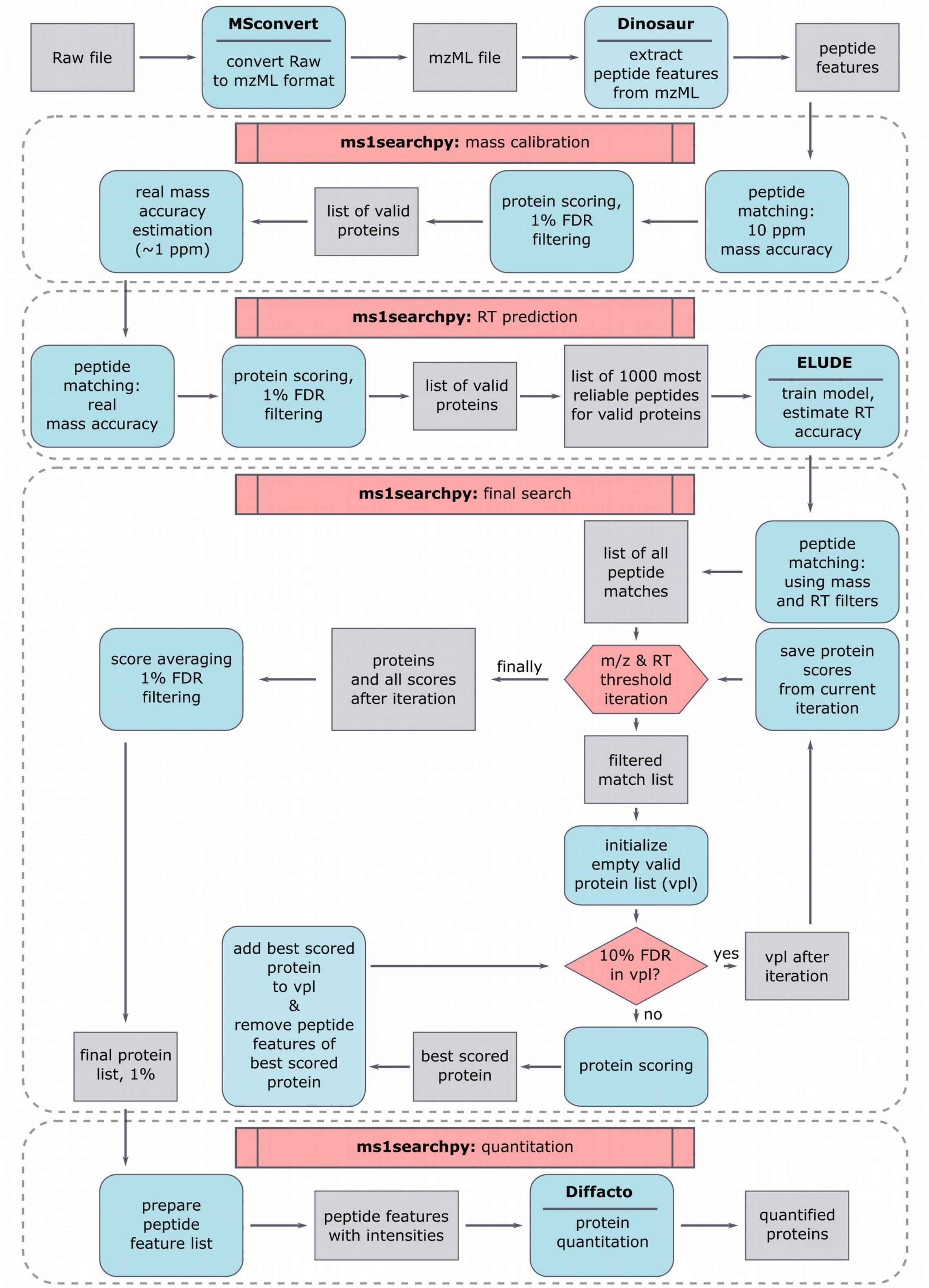


**Supplementary Figure S1.** Detailed workflow for ms1searchpy.

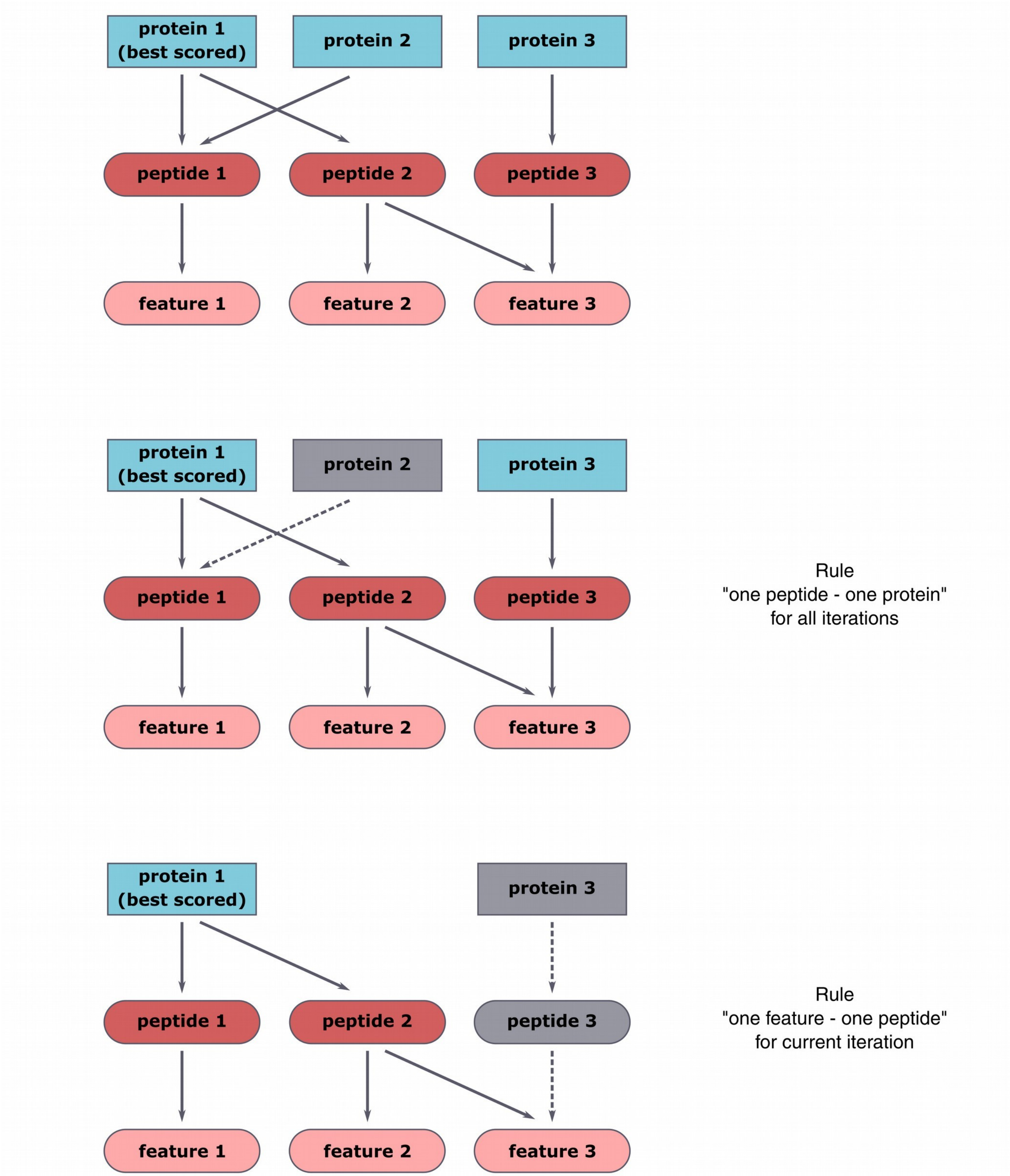


**Supplementary Figure S2.** Scheme of establishing relationships between the features in MS1 spectra, the peptides, and the proteins. “One peptide - one protein” rule states that all shared peptides are used for scoring the best scored protein only obtained at the previous protein search iteration step. “One feature - one peptide” rule states that all features belonged to the best scored protein are excluded from the following iterations.

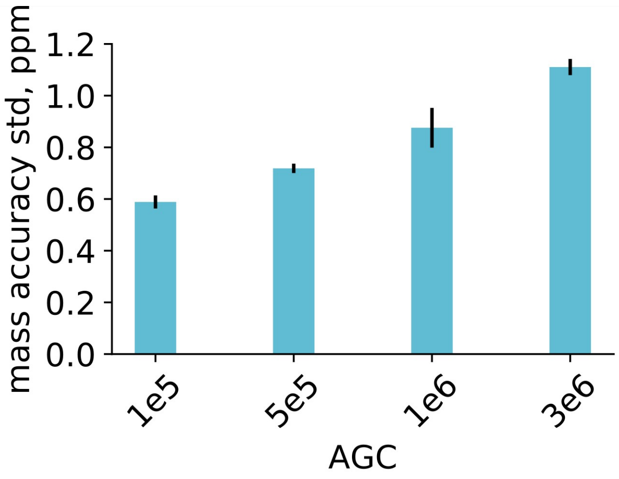


**Supplementary Figure S3.** Mass measurement accuracy for peptides of HeLa cell line using 5-minute HPLC separation gradient and different AGC settings in MS1 spectra. Results are shown for the average values obtained for 3 technical replicates
